# A Priori Activation of Apoptosis Pathways of Tumor (AAAPT) Technology 1: Sensitization of Tumor Cells Using Targeted and Cleavable Apoptosis Initiators in Gastric Cancer

**DOI:** 10.1101/492306

**Authors:** Wu Han, Li Yan, Raghu Pandurangi

**Affiliations:** Department of Oncology, Zhongnan Hospital of Wuhan University, Hubei Key Laboratory of Tumor Biological Behaviors and Hubei Cancer Clinical Study Center, Wuhan, China; Department of Peritoneal Cancer Surgery, Cancer Center of Beijing Shijitan Hospital, The Capital Medical University, Beijing, China; Sci-Engi-Medco Solutions Inc. and Amplexi-LLC, 573, Lexington Landing Pl, St Charles, MO 63303-1750

## Abstract

Cancer cells develop tactics to circumvent the interventions by desensitizing themselves to interventions. The principle route of desensitization includes activation of survival pathways (e.g. NF-kB, PARP) and downregulation of cell death pathways (e.g. CD95, ASK1). As a result, it requires high dose of therapy to induce cell death which, in turn damages normal cells through the collateral effects. Methods are needed to sensitize the low and non-responsive resistant tumor cells in order to evoke a better response from the current treatments. Current treatments including chemotherapy can induce cell death only in bulk cancer cells sparing low-responsive and resistant tumor cells. Here we report a novel tumor sensitizer derived from the natural Vitamin E analogue (AMP-001). The drug design is based on a novel “A priori activation of apoptosis pathways of tumor technology (AAAPT) which is designed to activate cell death pathways and inhibit survival pathways simultaneously. It involves an inbuilt targeting vector which targets tumor specific Cathepsin B, overexpressed by many cancers including gastric cancer. Our results indicate that AMP-001 sensitizes gastric cancer cells which resulted in expanding the therapeutic index of front-line chemotherapy doxorubicin both *in vitro* and *in vivo* nude mouse model. The synergy between AMP-001 and doxorubicin could pave a new pathway to use AMP-001 as a neoadjuvant to chemotherapy to achieve a better efficacy and reduced off-target toxicity.

## Introduction

Tumor cells have a remarkable ability to circumvent endogenous and exogenous toxicities by deactivating cell death pathways and thereby desensitizing themselves to interventions^1^. Defects in apoptosis pathways (e.g. CD95, APO-1/Fas) would make tumor cells insensitive to chemotherapy^2^. For example, loss of CD95 signaling pathway resulted in the development of cancer^3^ and resistance to chemotherapy^4^. Restoration of the apoptotic machinery with apoptosis inducing ligands (apoptogens) is an area of active investigation in the drug development. Several agents have been developed to activate TRAIL^5^, to downregulate Bcl-2^6^ and to restore function of mutated p53 in order to sensitize tumor cells. Similarly, mitogen activated protein kinase MAPK signaling pathways are also involved in desensitization of tumor cells. The advantage of sensitizing tumor technology is to evoke a better response from the existing treatments in terms of reducing the therapeutic dose without compromising efficacy and a potential reduction of dose related off-target toxicity. Consequently, tumor sensitizing agents, potentially can be used as neoadjuvant to chemotherapy to expand the therapeutic index of chemotherapy^7^.

Tumor sensitivity to chemotherapy *in vivo* is shown to be dependent on spontaneous baseline tumor apoptosis index^8^. Aggressive or incurable cancers show low tumor apoptosis index (AI) and low sensitivity to chemotherapy compared to benign cancer^9^. In fact, the spontaneous levels of apoptosis are strong predictor of treatment response^10^. For example, low baseline apoptosis index (AI) of patient tumors (> 67%, see Fig 4 in ref 3g), is directly correlated to non-respondent patients to chemotherapy. Lower the baseline apoptosis index of tumor, least the response from chemotherapy and vice versa (Fig 3 in ref 3h). The overall 5-year survival rates for the patient group with high AI (> 0.97) and low AI (< 0.97, p =0.001) were 89.6 % and 69.2 % respectively^11^ indicating that AI could be a potential biomarker of risk stratification of tumors/patients to see who responds better to which treatments.

We have developed a novel technology, “A priori Activation of Apoptosis Pathways of Tumor” (AAAPT) as a strategy to sensitize low responsive tumor cells in order to evoke a better response from chemotherapy^12^. The goal is to understand the molecular biology of desensitization tactics of tumor cells to bypass the intervention by reactivating specific dysregulated apoptosis pathways selectively in tumor sparing normal cells. Since ubiquitous or systemic activation of apoptosis can induce undesirable neurodegeneration and myelosupression^13^, targeting is essential^14^.

Gastric cancer remains the second leading cause of cancer-related deaths in the world^15a^. Almost two-thirds of new diagnoses occur in developing countries with 42% in China alone, remaining high in developed countries^15b^. Although, surgery is the main treatment for regressing gastric cancer, the majority patients first diagnosed with gastric cancer have already presented local or distant metastasis. Thus, adjuvant or perioperative chemotherapy and molecule-targeted chemotherapy have been prescribed for gastric cancer, due to their proved benefits in controlling metastasis, reducing cancer recurrence and increasing overall survival^16^. Gastric cancer cells are also known to downregulate CD95 pathway which makes them insensitive to chemotherapy^17^. Hence, we investigate the tumor sensitizing potential of a leading AAAPT molecule AMP-001 in gastric cancer to evoke a better response or synergistic effect with a standard chemotherapy for example, doxorubicin. doxorubicin (dox) is a first-line anticancer agent which is widely used in clinical therapeutic regimens for a variety of cancers including gastric cancer. Nevertheless, the clinical application is limited due to the drug resistance and adverse effects, including cardiomyopathy, typhilitis, and acute myelotoxicity^18^. To decrease dox-induced dose-dependent toxicity, enhancing the effective therapeutic dox dose via combination with nontoxic or selective apoptotic inducing ligands has been investigated^19^. Here, we propose novel targeted tumor sensitizers which addresses how the cancer cells circumvent interventions by reactivating cell death pathways. Since desensitization tactics by cancer cells are successful in thwarting the efficacy of treatments, irrespective of nature of interventions (targets), a fundamental reversal of cancer cells tactics may have high impact on several modes of interventions (e.g. chemotherapy, radiation and immunotherapy). Here, we report to demonstrate the synergistic effect of AMP-001with doxorubicin *in vitro* and *in vivo* in a gastric cancer animal model.

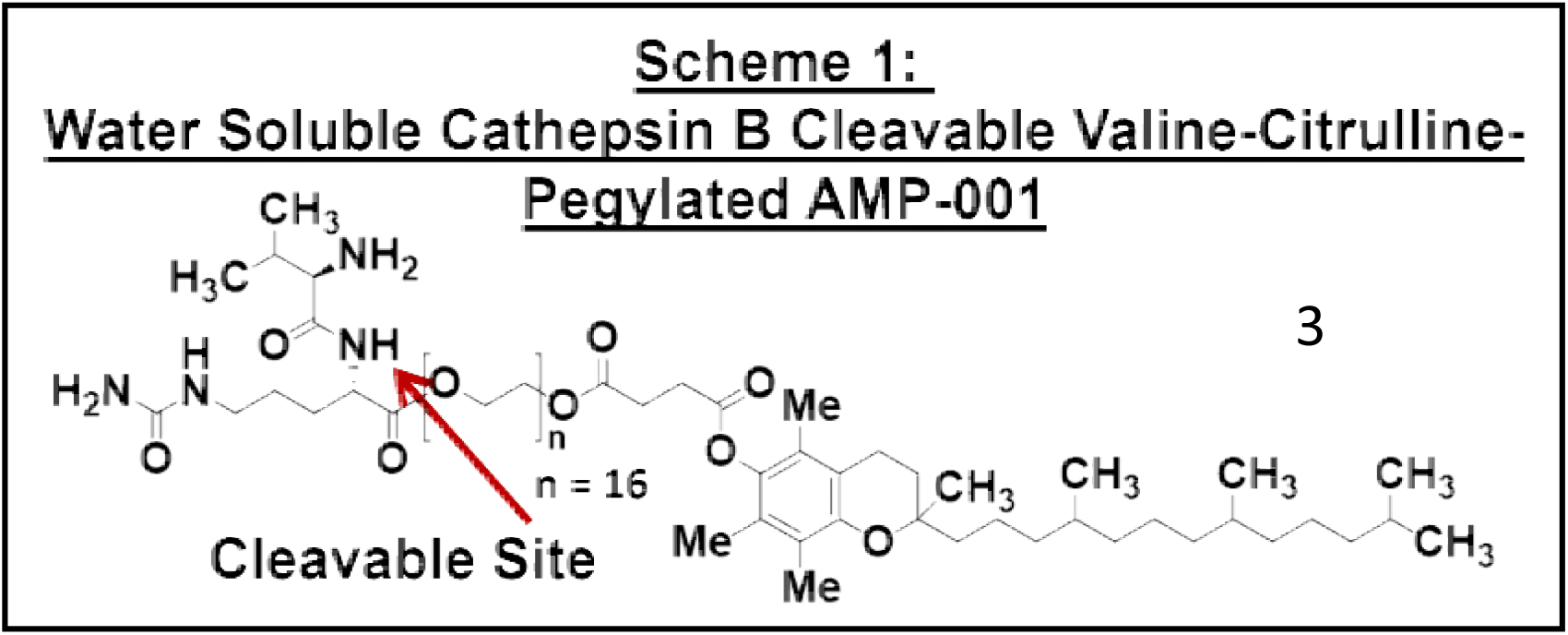

### Rational Drug Design

The drug design involves pegylation of α-tocopheryl succinate (apoptogen) with a dipeptide linker valine-citrulline which is cleavable by tumor specific Cathepsin B enzyme giving rise to a final molecule AMP-001(Scheme 1). This design provided impetus for a) targeting apoptogen to cancer cells using tumor specific biomarker Cathepsin B sparing normal cells, b) releasing the drug near tumor sites through the release of drug by cleaving at valine-citrulline link, c) pegylation to make it water soluble, d) enhancing the bioavailability of AMP-001and e) to keep it intact as a pro-drug in blood circulation for a long time to reach tumor sites. The rationale behind using cathepsin B cleavable linkers is based on the better safety profile of Cathepsin B cleavable prodrugs doxorubicin^20a^, paclitaxel^20b^ and antibodies^20c^ (Seattle Genetics) compared to unmodified drugs. Cathepsin B is known to a) be a tumor specific biomarker and b) cleave valine-citrulline substrate^21^. For example, high delineation of tumor from the surrounding tissues by cathepsin B sensitive optical probes^22^ shows that this enzyme is restricted to invading cancers. Cathepsin B cleaves AMP-001 at the citrulline-OH bond to release pegylated apoptogen at tumor sites.

Previous studies suggested that α-tocopheryl succinate could destroy mitochondrial function and cause the oxidative damage, resulting in over production of reactive oxygen species (ROS) and apoptosis. Although, parent molecule α-tocopheryl succinate (α-TOS) showed promising anti-tumor properties *in vitro*, studies in an immunocompetent mouse *in vivo* model showed that α-TOS was not only ineffective at the published doses, but resulted in severe side effects due to lack of targeting^23^. Elevated ROS levels can render cancer cells more sensitive to agents that cause oxidative stress^24-25^. There are many anti-cancer drugs such as daunorubicin^26a^, cisplatin^26b^, paclitaxel^26c^ and trisenox^26d^ that can augment ROS production in cancer cells, despite their inability to target cancer cells leading to non-specific side effects.

Here, we examined the effect of combining AMP-001 and doxorubicin for a potential synergistic effect in human gastric cancer both *in vitro* and *in vivo*. These results indicate that low dose pretreatment of tumor *in vivo* prepares tumor to make it sensitive to doxorubicin by showing a cumulative and significant tumor regression compared to individual drugs. A potential mechanism of action is being suggested based on the limited data.

## Materials and Methods

### Cell Culture and Treatment

The human gastric cancer cell lines MGC-803 and SGC-7901 were cultured in Dulbecco’s modified Eagle’s medium (DMEM) supplemented with 10% FBS, 100 IU/ml penicillin, 100 μg/ml streptomycin in a humidified atmosphere with 5% CO_2_ at 37°C.

### Reagents

The novel compound AMP-001 was designed and synthesized by Sci-Engi-Medco Solutions Inc (SEMCO). AMP-001 was dissolved in distilled water and stored at -20°C until use. The drug doxorubicin was purchased from Sigma-Aldrich Biotechnology. For all AMP-001/D0X combinational treatment, cancer cells were pretreated with AMP-001 before doxorubicin was added into the culture. All above micromolecular reagents were dissolved and saved according to manufacturer specifications.

### Cell Survival and Growth Assay

The detection of cell viability and cell growth were performed with cholecystokinin-8 (CCK-8) assay (Dojindo, Japan). Briefly, 6000 viable gastric cancer cells were seeded in 96-well plates. After specific treatment, each well was mixed with 10 μl CCK-8 and incubated for additional 1h. The OD values were detected at an absorbance of 450nm.

### Flow Cytometry Analysis

MGC-803 and SGC-7901 were placed in 12 well plates overnight, and then treated with AMP-001. Cells were then harvested, washed twice with pre-cold PBS, and evaluated for apoptosis by double staining with FITC conjugated Annexin V and Propidium Iodide (PI) kit (Multi Sciences) for 30 min in dark. For cell cycle, harvested cells were labeled with PI (5 mg/ml) in the presence of binding buffer (Multi Sciences) in darkness for 30 min. For JC-1 assay, the treated cells were incubated with JC-1 (Beyotime Institute of Biotechnology, Nanjing, China) for 30min at 37°C, and then washed twice with PBS. Beckman flow cytometer was used for final detection. Flow Jo vX.0.7 software was used to analyze the data.

### Measurement of intracellular ROS level

The intracellular ROS level was measured by flow cytometry as described previously^24^. Briefly, 2×10^5^ cells were placed in 12 well plates, allowed to attach overnight, and exposed to the treatment needed. Cells were stained with 10μM DCFH-DA (Sigma) at 37°C for 30 min with serum free culture. Cells were washed twice with pre-cold PBS and photographed or collected for fluorescence analysis using Beckman flow cytometer. In some experiments, cells were pretreated with 5mM NAC for 2h prior exposure to compounds and analyzed for ROS generation.

### Cell Mitochondria Isolation

Fractionation of intracellular mitochondria was performed with Cell Mitochondria Isolation Kit (Beyotime Institute of Biotechnology, Nanjing, China), according to the manufacturer’s protocols. First, 1×10^7^ cells were collected and homogenized. Density gradient centrifugation at 600g for 10min was applied to remove cellular debris, and supernatants were centrifuged at 11000g for 10min at 4°C to separate mitochondrial fractions. Then, supernatants containing cytoplasmic proteins were saved and the obtained mitochondria pellet was lysated for further immunoblotting.

### Western Blotting

Cells were washed with cold PBS twice and prepared in RIPA lysis buffer, and western blot analysis was performed as described previously[25]. Specific primary antibodies used were as follows: anti-Bcl-2 (50E3), Caspase-9 (C9), Phospho-p38 MAPK (Thr180/Tyr182), p38 MAPK, Phospho-p44/42 (Erk1/2) (Thr202/Tyr204), p44/42 MAPK (Erk1/2), Phospho-SAPK/JNK (Thr183/Tyr185), JNK2 antibodies were purchased from Cell Signaling Technology (USA). Anti-Bax antibody was purchased from Abcam. Anti-Cytochrome C and COX IV were purchased from Beyotime Biotechnology. Antibody against PARP1 was obtained from Proteintech and Caspase-3 from ABclonal Technology. Anti-GAPDH and β-ACTIN were purchased from Aspen (China). After incubating with the fluorescein-conjugated secondary antibody, the membranes were detected using an Odyssey fluorescence scanner (Li-Cor, Lincoln, NE).

### Transmission Electron Microscopy (TEM)

The treated cells were fixed in ice-cold 2% glutaraldehyde, scraped from the plates and post-fixed in 1% osmium tetroxide with 0.1% potassium ferricyanide, dehydrated through a graded series of ethanol (30%-100%), and embedded in Epon-812 monomer and polymerized. Ultrathin sections were cut with a diamond knife mounted in a Reichart ultramicrotome, contrasted with uranyl acetate and lead citrate, and examined in a Hitachi HT7700 transmission electron microscope operated at 120 kV^26^.

### In Vivo Xenograft Assay

Six-week-old athymic BALB/cA nu/nu female mice were purchased from Weitonglihua Laboratory (Beijing, China) and maintained in an Animal Biosafety Level 3 Laboratory at the Animal Experimental Center of Wuhan University. All animal experiments were performed according to the Wuhan University Animal Care Facility and National Institutes of Health guidelines. Approximately 5×10^6^ MGC-803 cells were harvested and suspended in 200μl of PBS and Matrigel (BD Bio-science) (1:1) and injected subcutaneously into the right flank of each mouse. After two weeks xenotransplantation, mice were respectively randomized into four groups and treated as follow: AMP-001 (10mg/kg i.p. every other day for 3weeks), DOX (1mg.kg i.p. every other day for 3 weeks), their combination, or saline as untreated vehicle. The size of subcutaneous tumors and mice weight were recorded every two days. The tumor volume (V) was calculated according to the formula V=0.5×l×w^2^, where l is the greatest diameter and w is the diameter at the point perpendicular to l. At the end of treatment, mice were sacrificed, and the tumors were removed and used for immunohistochemical staining.

### Immunohistochemical Staining

The xenograft tumor slides were incubated with the following primary antibodies: anti-CD31 was purchased from ABclonal and anti-Ki67 from Cell Signaling Technology (USA). Anti-rabbit or anti-mouse peroxidase-conjugated secondary antibody (ABclonal) and diaminobenzidine colorimetric reagent solution purchased from Dako (Carpinteria, CA) were used. The staining processes were according to standard methods.

### TUNEL Assay

The level of tumor tissue apoptosis in vivo was determined using an TdT-mediated dUTP nick end labeling in situ apoptosis detection kit (Roche, USA) according to the manufacturer’s protocol. The pictures were photographed by fluorescence microscopy at a 200x magnification.

### Cardiotoxicity Assay

The data were collected from Ionic Transport Assays Inc. In brief, the adult human heart cell line was created by reprogramming an adult human fibroblast cell line by retroviral expression of the reprogramming factors *sox7, oct4, nanog*, and *lin28* using MMLV viral constructs. Cardiomyocytes were derived from this engineered stem cell clone line as follows. Stem cell aggregates were formed from single cells and cultured in suspension in medium containing zebrafish bFGF (basic fibroblast growth factor) and fetal bovine serum. Upon observation of beating cardiac aggregates, cultures were subjected to blasticidin selection at 25 ug/ml to enrich the cardiomyocyte population. Cardiomyocyte aggregate cultures were maintained in Dulbecco’s modified Eagle’s medium (DMEM) containing 10% fetal bovine serum during cardiomyocyte selection through the duration of the culture prior to cryopreservation. At 30 to 32 days of culture the enriched, stem cell-derived cardiomyocytes were subjected to enzymatic dissociation using 0.5% trypsin to obtain single cell suspensions of purified cardiomyocytes, which were >98% cardiac troponin-T (cTNT) positive. These cells (iCell^®^ Cardiomyocytes) were cryopreserved and stored in liquid nitrogen before delivery to Ionic Transport Assays from Cellular Dynamics International, Madison, WI.

Cells were plated into 6 well plates that percolated with 0.1% gelatin. This was defined as culture day 1 for the purpose of this study. Cell plating media was changed at day 3 to cell maintenance media and cell maintenance media subsequently was changed three times a week. Day 5-7 cells were re-suspended with trypsin and re-plated as desired density (>10,000) at 96 well plate which percolated with 0.1% gelatin.

### Sample Preparation

3.75 mg AMP-001 was dissolved into 0.25 ml water to create a 10 mM stock solution. This stock solution was added to maintenance medium for a final concentration 500 μM which was then diluted to 200 μM, 100 μM, 10 μM and 1 μM AMP-001 solution. 3 mg Dox (Tocris, Cat No. 2252, Cas No.25316-40-9, MW=579.99) was dissolved in 0.5 ml water to create a 10 mM stock solution. This stock was stored in desiccators at room temperature. This stock solution was added to maintenance medium with 200 μM AMP-001 for a final concentration 20 μM which was then diluted to, 10 μM, 1 μM, 0.1 μM and 0.01 μM Dox solution. The CCK-8 kit used in these experiments to determine IC-50 values is a nontoxic, highly sensitive colorimetric assay for the determination of cell viability in cell proliferation and cytotoxicity assays. Raw data were measured on a Spectra Max micro-96 well plate reader and plotted using Prism 5. Transformation, normalization and nonlinear regression were used to analyze data. According best-fit values were used to obtain IC-50. DMSO concentrations less than 1% had no effect on cell viability

### Statistical Analysis

All experiments were performed at least three times. Data are presented as means ± SD. All statistical analyses were performed using GraphPad Prism 6.0 (GraphPad, SanDiego, CA). One-way ANONVA and Student’s t-test were applied to determine statistical significance. A value of p<0.05 was considered statistically significant.

## Results

### 1.0 Synthesis of AMP-001

The general synthesis of cathepsin B cleavable peptide conjugation with pegylated apoptogen and/or other apoptogens is accomplished through Fmoc chemistry to protect N end of peptide with Boc and then couple it with tocopherol derivatives using DCC in DMF. This was further cleaved by TFA and purified using HPLC method. Cathepsin cleavable compounds have been synthesized using the proprietary peptide synthesis technology. In brief, dipeptide valine-citrulline was synthesized using a microwave peptide synthesizer. The resin (containing 0.25 mmol of peptide anchors) was deprotected using piperidine resulting in the formation of the primary amine. The carboxylic acids of the Fmoc protected amino acids (1 mmol) were activated using COMU and conjugated to the primary amines of the growing peptide on the resin. The process of deprotection, activation and conjugation was repeated until the desired peptide was synthesized. Purification of the peptide linked pegylated apoptogen (AMP-001) was performed using a semi-preparative Kromasil C18, 5u column with a flow rate of 5.0 mL/min. HPLC solvents were 0.1 % TFA acetonitrile (solvent A) and 0.1% TFA in water (solvent B) to get off-white powder. The initial gradient A: B, t =0, 10: 90 and t = 30 100 % B. Analytical data: HPLC, R_f_ = 14.3, single peak with a purity 96.62 %. The compound is characterized using MS with a molecular ion peak at 1967 corresponding to M+H peak. The compound was soluble in water (100mg/mL).

### 2.0 Cytotoxicity of AMP-001 in Gastric Cancer Cell Lines

To investigate the cytotoxicity of AMP-001 on human gastric cancer cells, cell lines SGC-7901 and MGC-803 were treated with different concentration of AMP-001 *in vitro* for 48 hours and cell viability was detected using CCK-8 assay. As shown in Fig. 1A-B, AMP-001 inhibited the growth of SGC-7901 and MGC-803 cells respectively in a concentration dependent manner. The IC_50_ values of AMP-001 against SGC-7901 and MGC-803 were 14.25 and 16.91 μM, respectively (Fig 1C-D). Cell proliferation rates were also decreased in a dose and time dependent manner as shown in Fig. 1E-F respectively.

**Fig 1.**
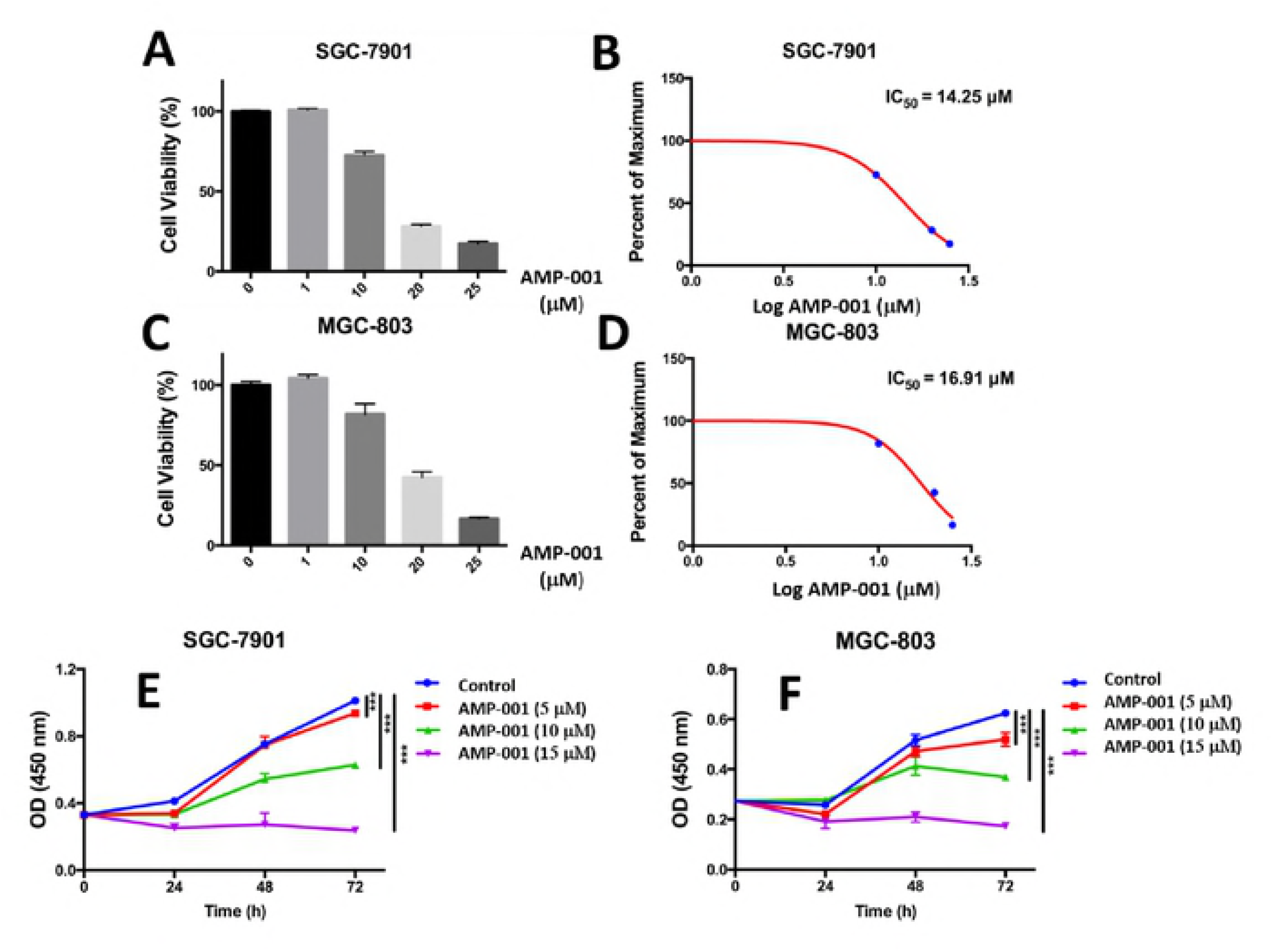
A& C: Viability assay for AMP-001 in Gastric cancer cells SGC-7901 and MGC-803 respectively, B&D: IC50 curves, E&F: Concentration and time dependent cell proliferation assay for AMP-001.

### 3.0 AMP-001 Induces Apoptosis and Potentiates Doxorubicin-Induced Apoptosis in Gastric Cancer Cells

The pro-apoptotic effect of AMP-001 was assessed using Annexin V/ staining assay. As shown in Figure 2, two gastric cancer cell lines SGC-7901 and MGC-803 showed significant apoptosis after 48h treatment with IC-50 dose of AMP-001. Although AMP-001 itself was able to induce cell apoptosis, the aim is to use it as a tumor sensitizer in conjunction with chemotherapy (e.g. doxorubicin) to see the potential synergy with it. This will enable us to extend the sensitizing potential of AMP-001 to other FDA approved chemotherapeutics used to treat other kinds of cancers. We, thus investigated whether the low dose of AMP-001 could be used in combination with doxorubicin, a widely used chemotherapeutic drug in the treatment of gastric cancer. MGC-803 and SGC-7901 cells were pre-incubated with or without AMP-001 (10 μM) for 8 hrs. and then treated with DOX (1 μM) for 48 hrs. before analyzed by FACS. Pre-incubation of gastric cancer cells with AMP-001 for 8 hours rendered cancer cells much more susceptible to doxorubicin induced cytotoxicity at a concentration at which AMP-001 itself did not cause substantial cell death. Fig 3A shows pictures of gastric cancer cells by transmission electron microscopy before and after AMP-001 treatment where the combination of doxorubicin and AMP-001 showed lower number of cancer cells compared to individual AMP-001 (10 μM) or doxorubicin (1 μM). This was confirmed by the flow-cytometric analysis of quantification of apoptosis, showing that AMP-001 enhanced early cellular damage induced by doxorubicin in MGC-803 and SGC-7901 cells by several folds (Fig 3B). In addition, combination treatment of cancer cells with 10μM AMP-001 significantly reduced the IC_50_ of doxorubicin in SGC-7901 and MGC-803 cells (~ 6.68 and 9.68 fold reduction, respectively) when compared with doxorubicin only treatment (Fig 4 A-B), thus, establishing the synergy between doxorubicin and AMP-001. These results will have a broader connotation that potentially, one can reduce the chemotherapeutic dose clinically without compromising efficacy and in the process, could reduce potentially dose related off-target toxicity. In other words, it is possible to expand the therapeutic index of chemotherapeutics when AMP-001 is used as a neoadjuvant to chemotherapeutics which are currently used for treating cancer.

**Fig 2:**
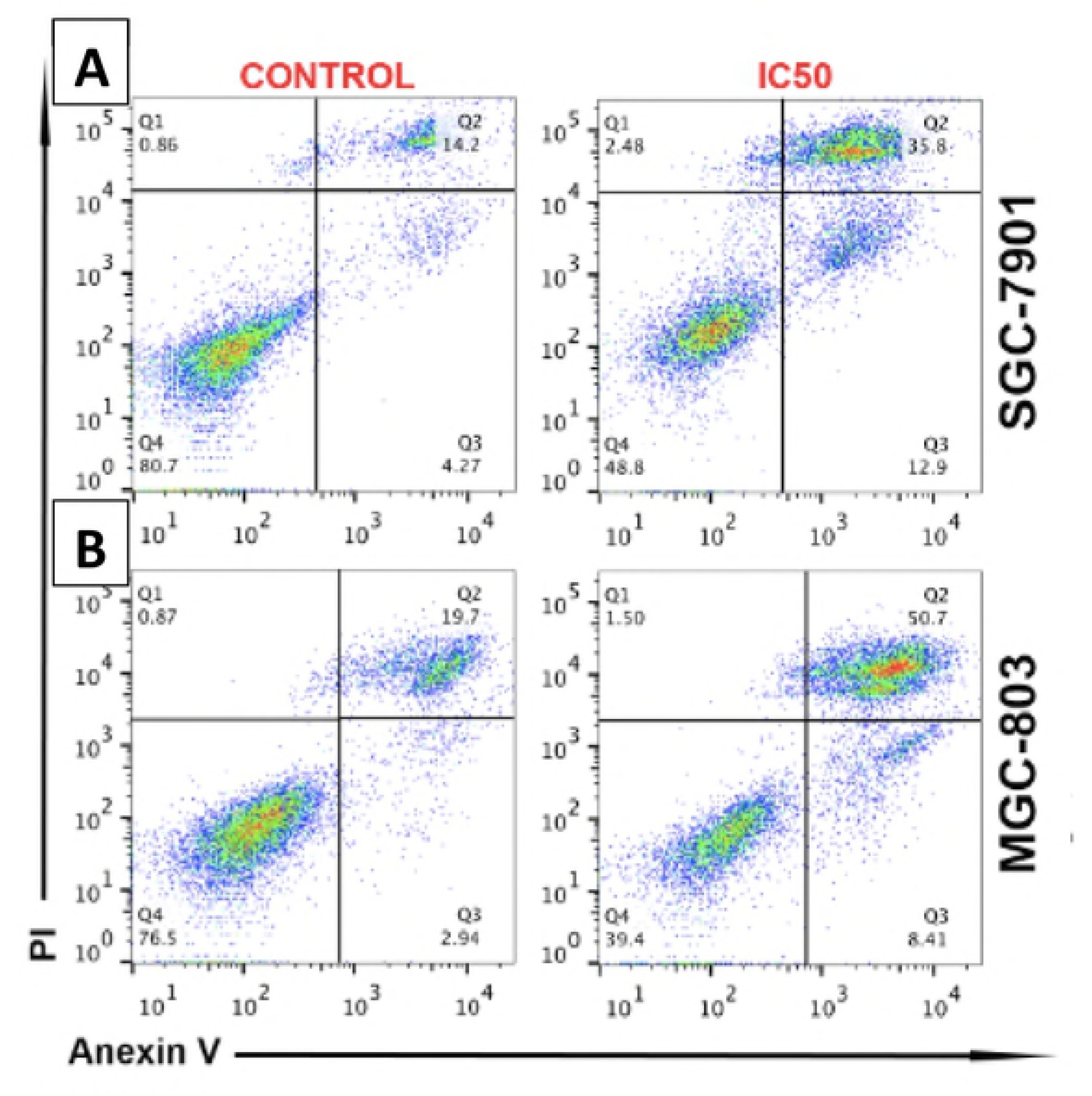
A-B: Pro-apoptotic Effect of IC-50 dose of AMP-001 in SGC-7901(14 μM) and MGC-803 (17μM) Gastric Cancer Cells Using Flow Cytometry with Annexin and PI Staining. AMP-001 increased cell death significantly in gastric cancer cells.

**Fig 3A:**
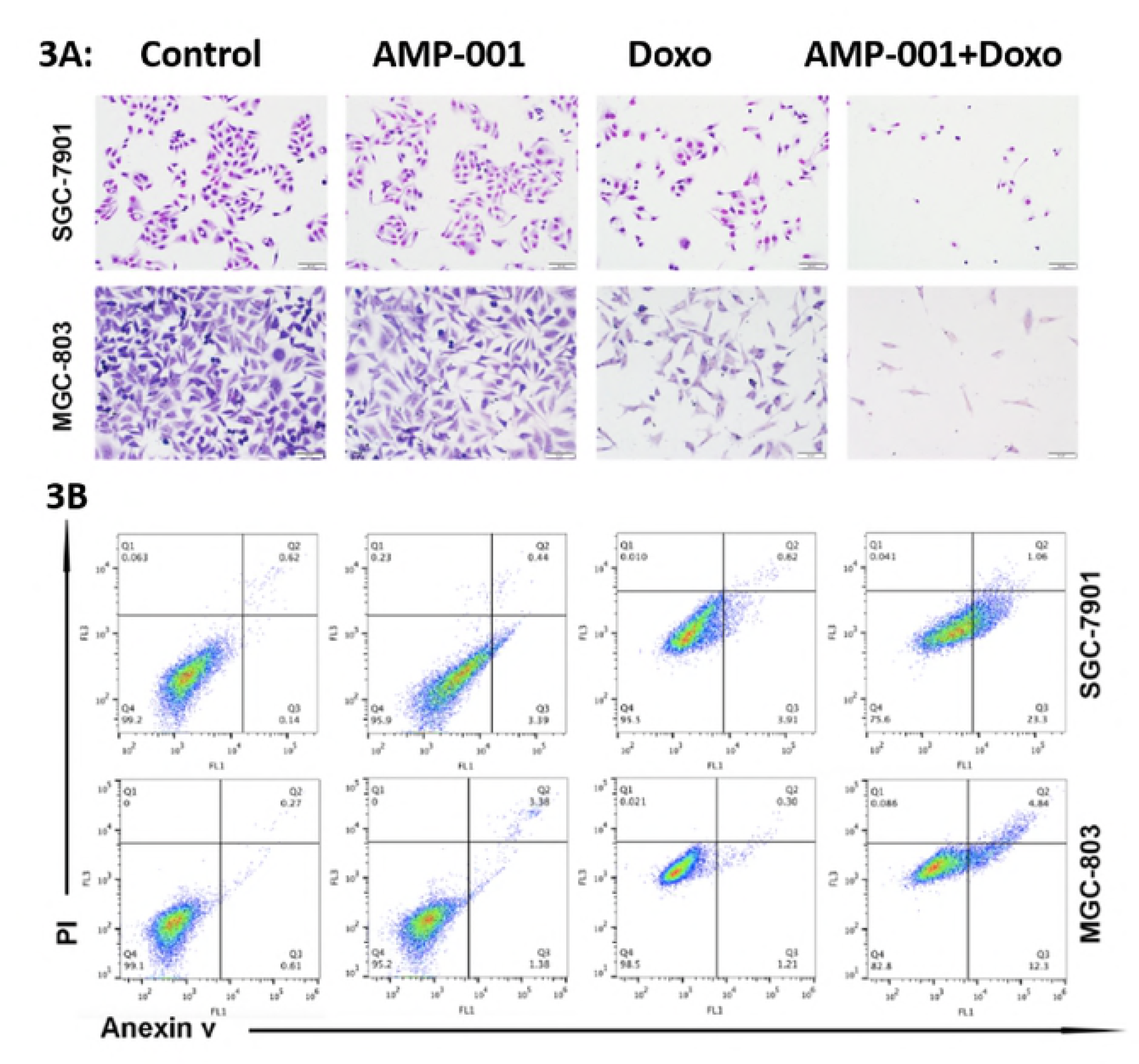
Synergy of AMP-001 (10 μM) with Doxorubicin (1 μM) in SGC-7901 and MGC-803 respectively, assessed through transmission microscopy. Combination of AMP-001 and Doxorubicin yielded cumulative cell death by an order of magnitude compared to individual drug treatments. 3B: Synergy of AMP-001 (10 μM) with Doxorubicin (1 μM) corroborated with FACS data. The combination of AMP-001 and doxorubicin showed higher cell death compared to individual drugs.

**Fig 4:**
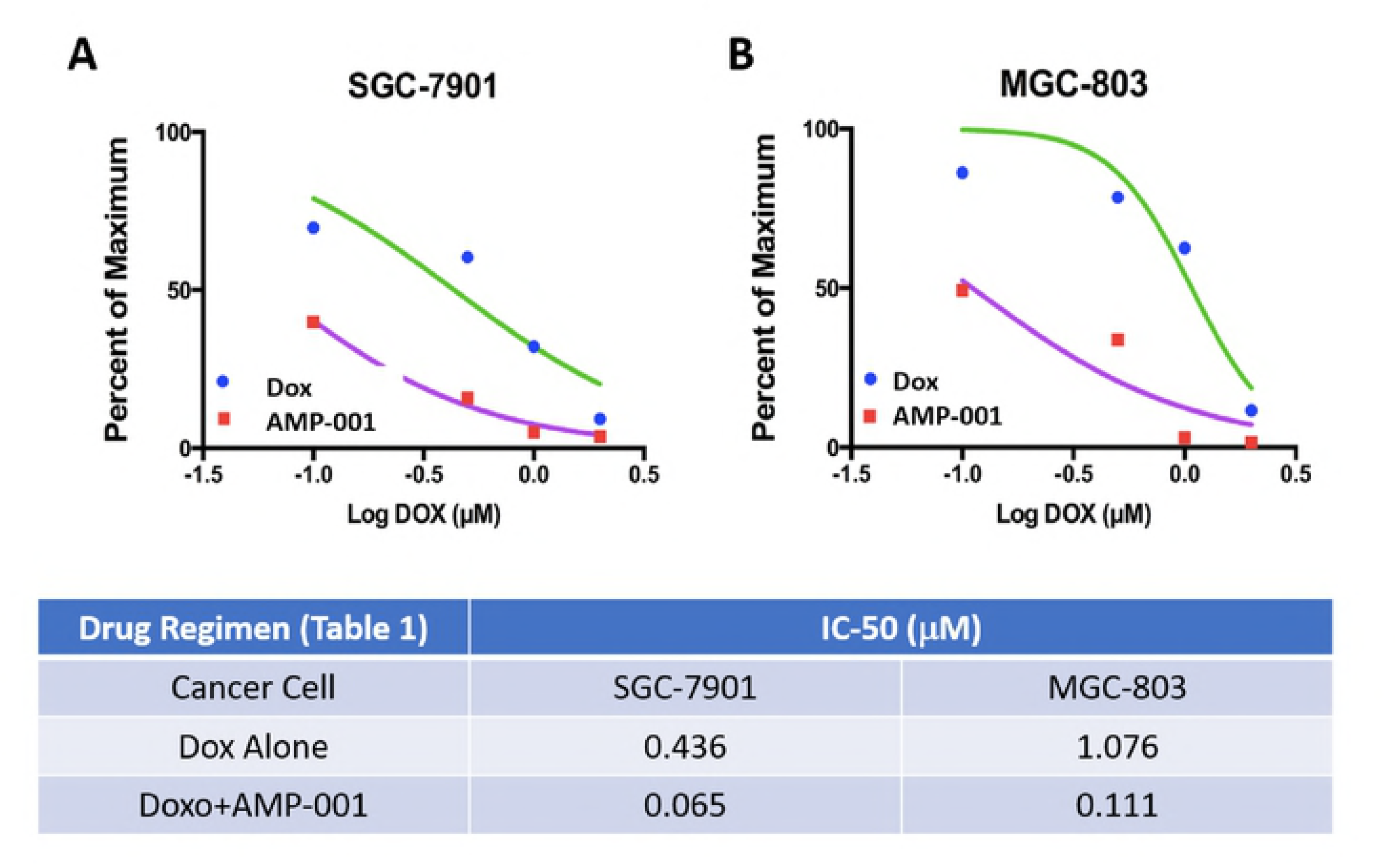
Quantitative Drift of IC_50_ value for the combination of AMP-001 and Doxorubicin in A) SGC-7901 and B) in MGC-803 cancer cells respectively.

### 4.0 AMP-001 Enhances DOX-induced Apoptosis through Mitochondrial Dysfunction

As AMP-001 could induce human gastric cancer cells apoptosis at a decent dose, the potential mechanism of cell death through reactive oxygen species (ROS) accumulation was investigated. As shown in Fig. 5 A-B, pretreatment with AMP-001 for 24h before doxorubicin treatment in SGC-7901 and MGC-803 cells caused a significant increase in DCF-reactive ROS than doxorubicin alone. It is well known that the excessive ROS generation causes oxidative stress and impairs membrane potential, resulting in mitochondrial dysfunction^28-29^. Hence, analysis of mitochondrial potential was carried out using polychromatic dye JC-1 (5,59,6,69-tetrachloro-1,19,3,39-tetraethylbenzimidoazolylcarbocyanino iodide) which forms red-orange clusters with high membrane potential mitochondria. while low membrane potential mitochondria show green staining. The results revealed a significant decrease in mitochondrial membrane potential through the enhanced monomer green fluorescence post combinational treatment (12.1%) compared to individuals AMP-001 (2.1%) and doxorubicin (5.7%) (Fig 5C).

**Fig 5:**
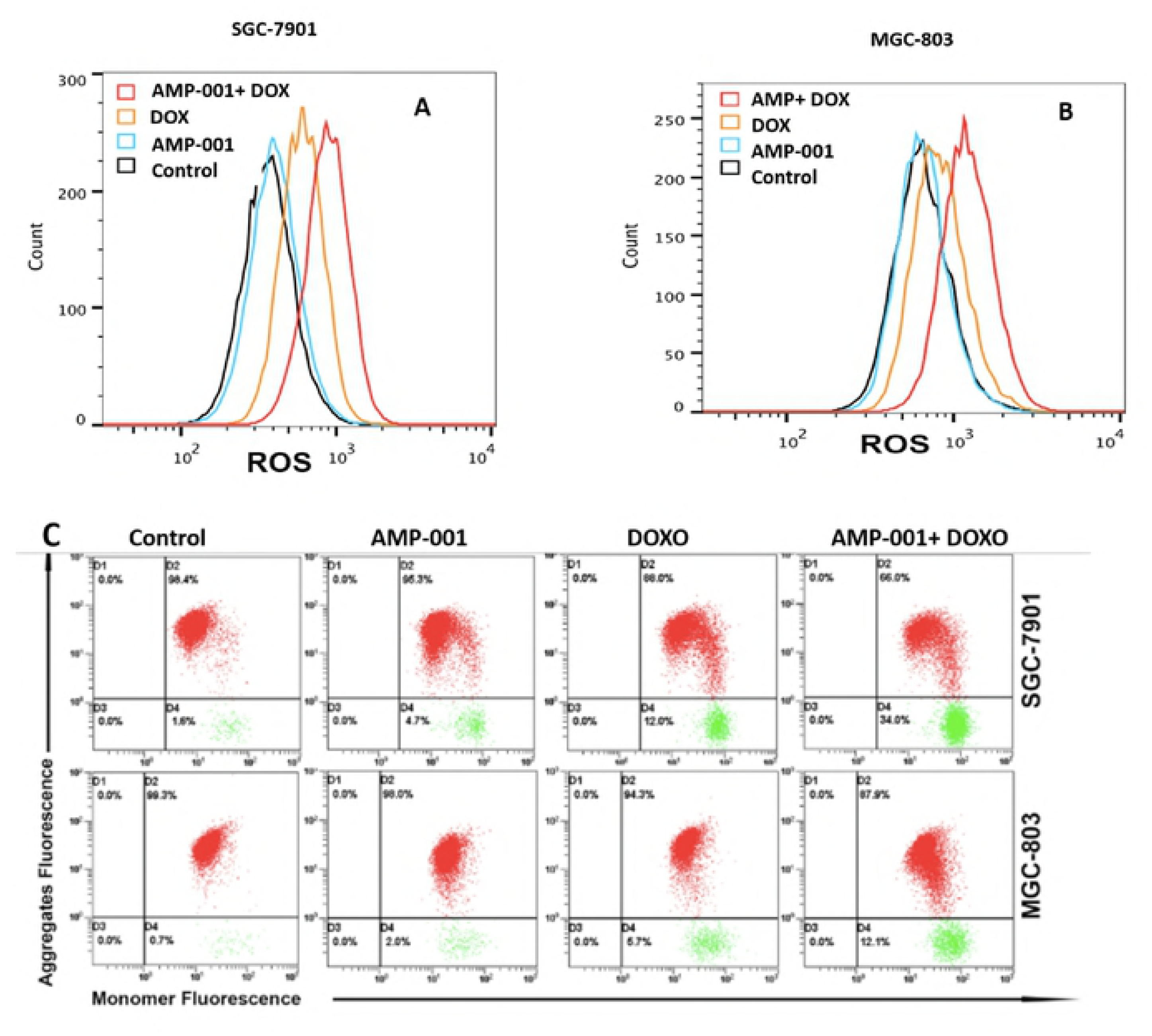
A-B: Enhancement of potentiation of reactive oxygen species (ROS) for the combination of AMP-001 and doxorubicin compared to individual drugs, C: Decrease in mitochondrial potential (enhancement of green fluorescence) using JC-1 dye in SGC-7 and MGC-803 cancer cells respectively.

### 5.0 AMP-001 and Doxorubicin Synergistically Induce Sustained Mitogen Activated Protein Kinase (MAPK) and CD 95 Pathways Activation

The mitogen activated protein kinase (MAPK) signaling plays a critical role in the outcome and the sensitivity to anticancer therapies. MAPK signaling is also associated with various cellular stress and stimuli and has been shown to contribute to induction of apoptosis^30^. P38 MAPK, c-jun N-terminal kinase (JNK) and extracellular-regulated kinase 1/2 (ERK1/2) are the three major kinases in MAPK family. We thus examined whether AMP-001 could enhance the apoptosis inducing effects of doxorubicin through MAPK signaling activation. The activated status of MAPKs, ERK and p-38MAPK was characterized using MAPK antibodies that recognize the dually phosphorylated peptide sequence representing the catalytic core of active MAPK enzymes. In all experiments, cells were preincubated with 10 μM of AMP-001for 8 hrs. before adding doxorubicin and analyzed post 24 hrs. The results revealed clear increases in p-p38 MAPK, p-JNK and p-ERK activity following the combination treatment compared with the control and with AMP-001 alone (Fig 6A). It is to be noted that AMP-001 and doxorubicin have incremental effect on these activities. However, in combination with doxorubicin, there is a significant enhancement in the intensity of the downstream phosphorylated bands. These results suggested that MAPK pathways could play important role in doxorubicin/AMP-001-induced apoptosis. For CD 95 pathway activation assessment, Fas resistant MDA-MB-231 TNBC cells were treated with different concentrations of AMP-001 (5, 10 and 50 μM) and assess the expression of CD95 (43 KDa) by Western blotting. Isolation of membrane proteins after cell lysis showed concentration dependent CD95 expression compared to untreated control (Fig. 6B). Translocation of CD95 from cytosol to the membrane is a common observation for CD95 Trail synergy^31^ which influences the cellular sensitivity to Fas death receptor pathway.

**Fig 6A:**
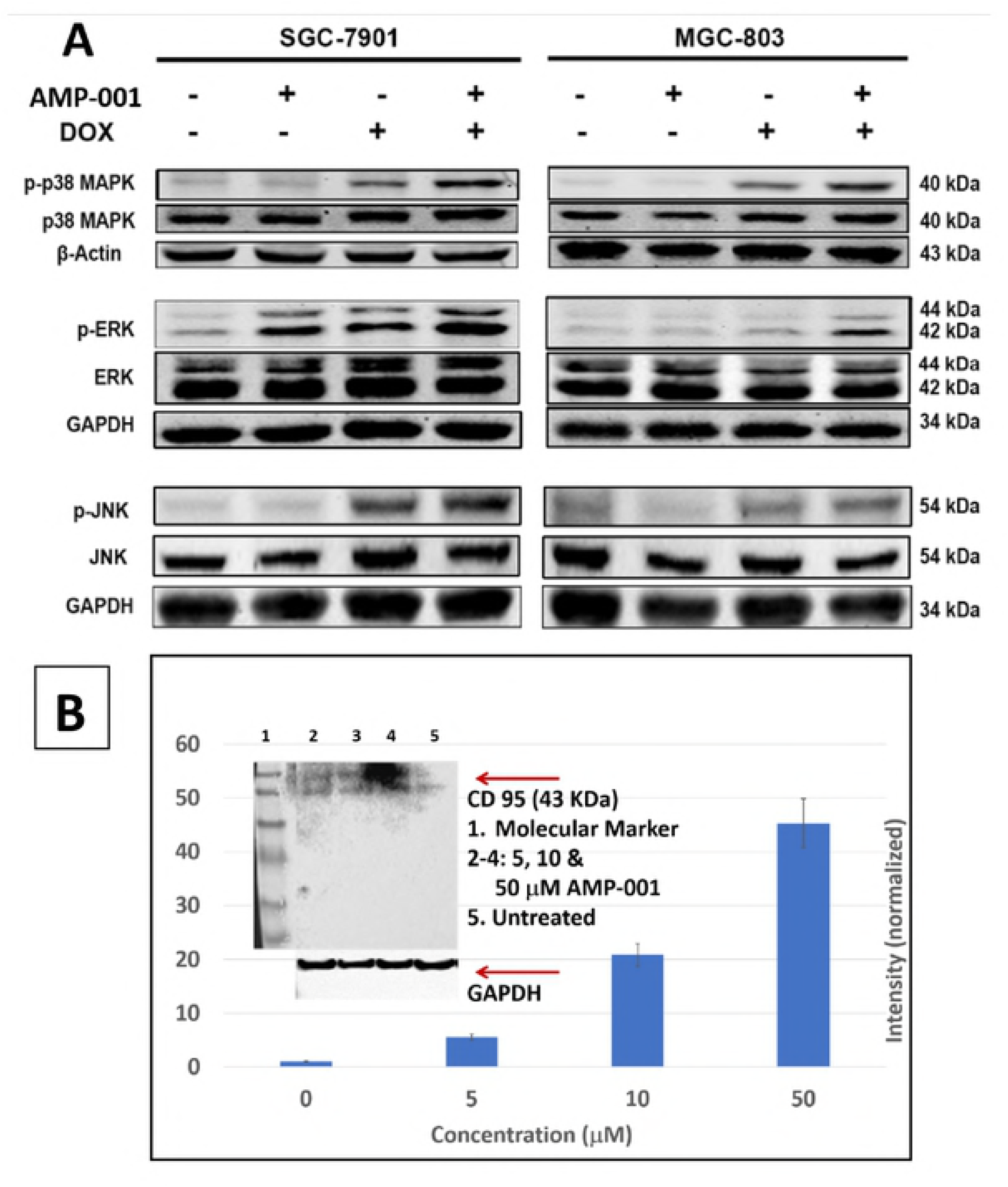
Potential Mechanism of Action for AMP-0012. Activation of MAPKs signaling pathway during AMP-001 and Doxorubicin combination treatment induced apoptosis, 6B: Expression of CD95 band in post treated triple negative breast cancer cells MDA-MB-231 by AMP-001. CD95 pathways is associated with the sensitization of tumor cells.

### 6.0 AMP-001 Amplifies the Therapeutic Effect of Dox in a Xenograft Gastric Cancer Model in Nude Mice

The sensitizing potential of AMP-001 has paved the way for evoking a better response from chemotherapy, particularly doxorubicin *in vitro*. Hence, it is may be essential if the same thing is true in an *in vivo* situation. Doxorubicin either alone or in combination with AMP-001 were injected i.p in a xenograft gastric cancer model in nude mice. MGC-803 cells were implanted subcutaneously in the right flank of nude mice. After a week, the mice were randomized into 4 groups and treated as described in the experimental protocol. Animals were sacrificed after 3 weeks treatment. We found that low dose (10 mg/Kg) AMP-001 alone did not inhibit the growth of the tumor at that dose compared to control, while doxorubicin alone (1mg/Kg) inhibited tumor growing to a certain degree. However, the combination of AMP-001 and doxorubicin was more effective in reducing tumor burden (Fig 7A). The tumor volume and tumor weight in the combinational group was significantly lower than either doxorubicin or AMP-001. (Fig 7B, p < 0.001). Additionally, all mice did not undergo obvious body weight loss or abnormal symptoms in drug treatments (Fig 7D). In alternate studies on AMP-001 in triple negative breast cancer model, we have shown that high dose of AMP-001 (~200 mg/Kg) alone can have significant tumor regression with no observable toxicity, despite higher dose^12^. Similarly, in the current studies also, there was no evident histopathological abnormalities were observed in the vital organs such as heart, liver and kidney estimated by HE staining (Fig 8). Ki67 (marker for cell proliferation) and CD31 (marker for micro vessel density) were also examined by immunohistochemistry in the excised tumor sections. Although, doxorubicin also downregulated the Ki67 expression, the combination group was most effective similar to the tumor regression studies. The trend of CD31 expression was consistent with Ki67 (Fig 8). In order to determine whether AMP-001/Dox combination treatment effectively promotes apoptosis in tumors, sections of excised tumors were also subjected to TUNEL assay. The results showed AMP-001 together with doxorubicin treatment had more TUNEL positive cells compared to either doxorubicin or AMP-001 (Fig 8). In summary, the combination of doxorubicin and AMP-001 yielded a superior response compared to either AMP-001 or doxorubicin in xenograft gastric cancer animal model.

**Fig 7:**
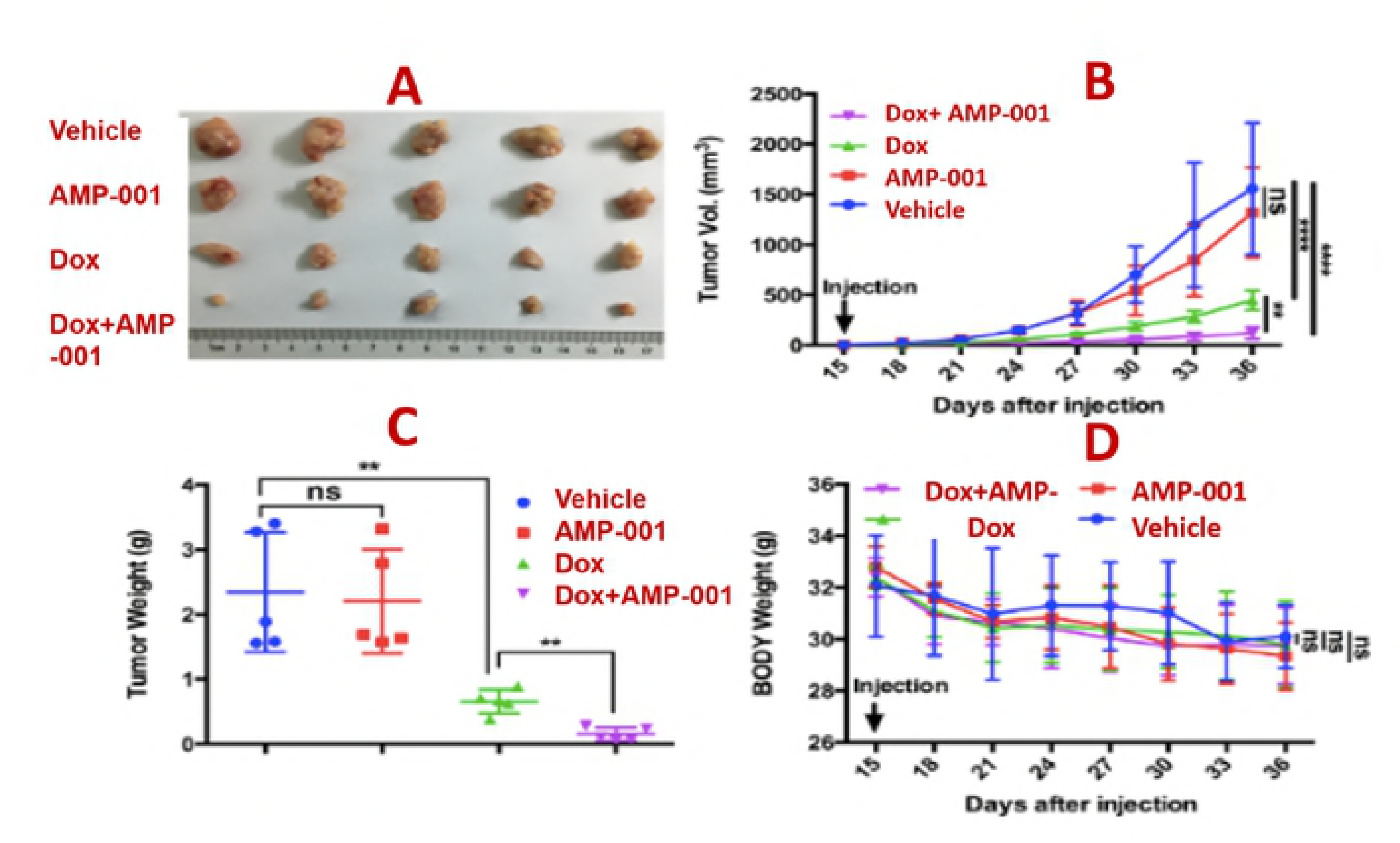
Synergy betwwen AMP-001 and Doxorubicin in xenograft gastric cancer nude mouse model: A: Dose dependent tumor size for untreated control, AMP-001, Doxorubicine and Combination of AMP-001 and Doxorubicin, B: Classic tumor V curve for tumor regression, C: Statistical tumor weight reduction comparison, D: Mouse body weight measurements post treatment.

**Fig 8:**
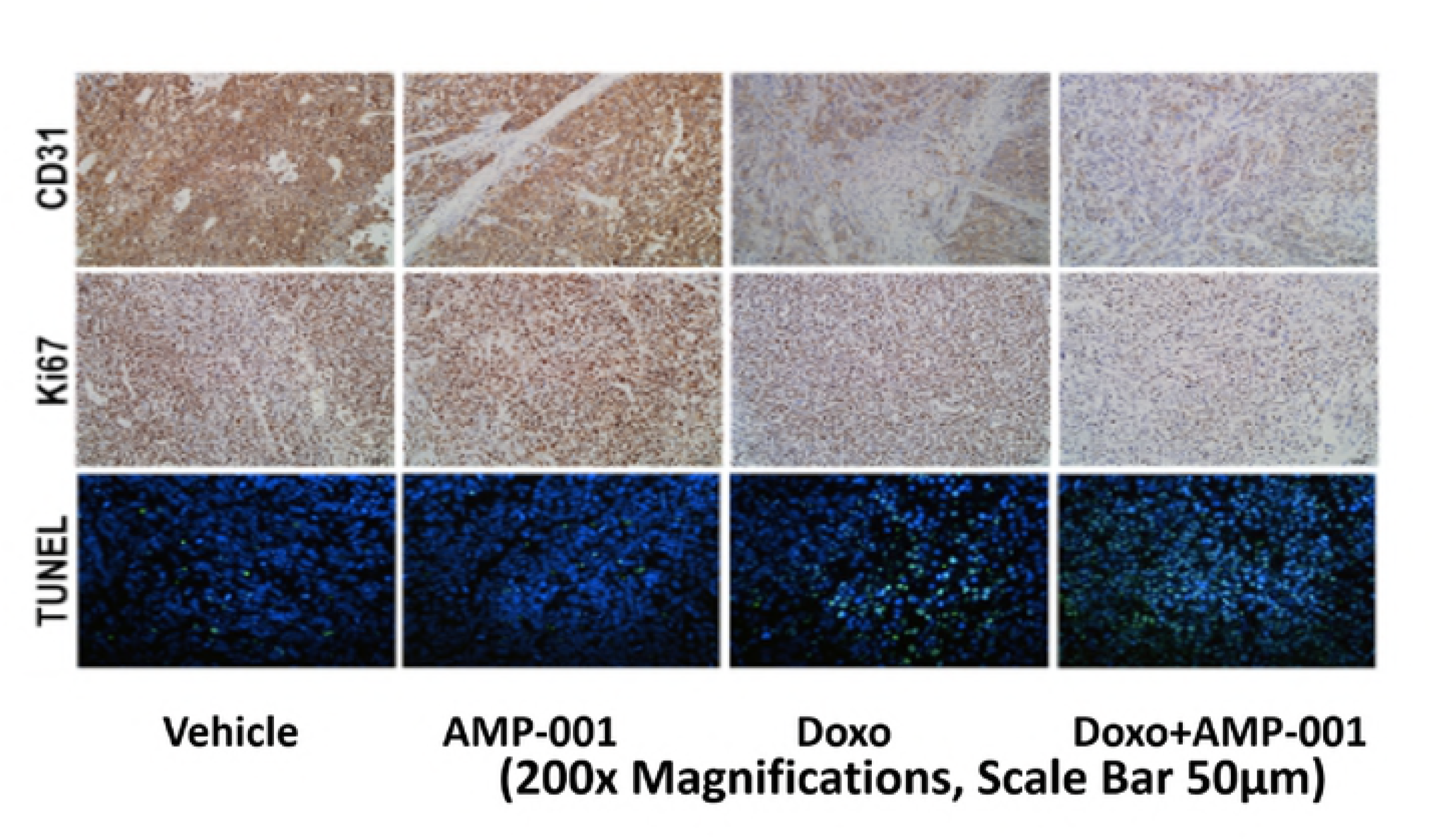
Representative immunohistochemical analysis of CD31, Ki67 and apoptosis in tumor sections Ex Vivo by TUNEL staining by combination of AMP-001 and Doxorubicin

### 7.0 Cardiotoxicity of AMP-001 Compared to Dox

The photomicrograph of cells in control experiment when the iPSCs are treated with 0.8% DMSO (Fig 9A) showed normal, beating cardiomyocytes. The treatment of the same cells with 50 0r 100 mM of AMP-001 did not change the morphology of cardiomyocytes, while 10 mM of dox induced significant cell death, shown by the disintegration of cardiomyocytes (Fig 9C). This pattern of normal beating for cardiomyocytes was seen at all concentrations including 250 μM AMP-001 (Fig 9D). AMP-001 showed IC-50 greater than 250 μM, compared to dox at 9.6 μM (25 times less, Fig 9F) which indicates a better safety profile for AMP-001 Even FDA approved drug (Sorafinib ~ 40 μM) has lower IC-50 compared to AMP-001 (Fig 9E). It is to be noted that IC-50 for cardiomyocytes has to be high (measure of safety) in contrast with IC-50 for cancer cells which should be low (measure of potency). AMP-001 also showed a high potency for cancer cells since IC-50 for MDA-MB-231 triple negative breast cancer cells to be 32 μM (Fig 9E). In essence, AMP-001 could be a potent antitumor agent with a better safety profile compared to dox.

**Fig 9.**
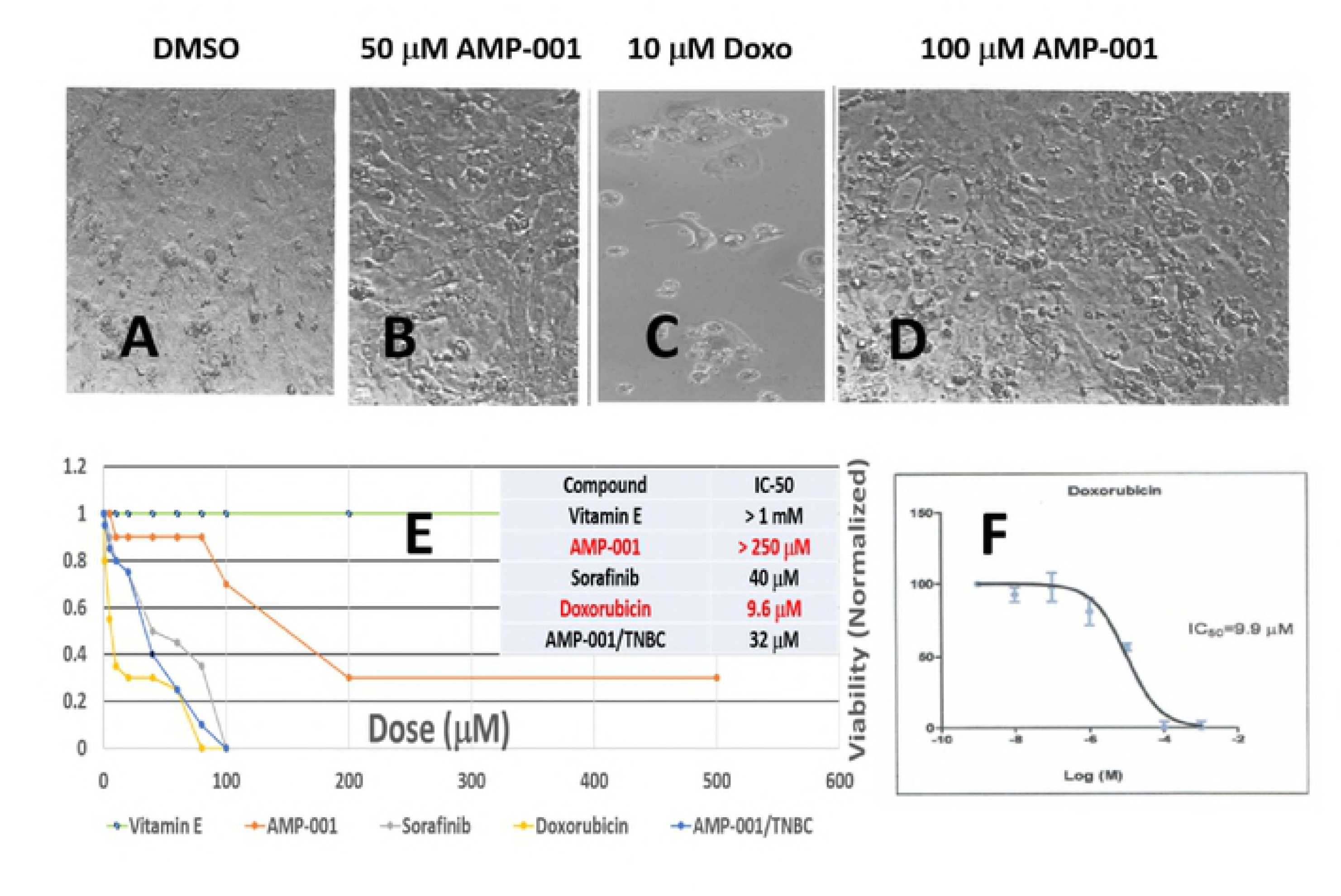
Photomicrographs of treated with A) control DMSO, B) 50 □M AMP-001, C) 10 μM Dox, D) 100 μM AMP-001 and E) IC-50 measurements for Vitamin E, AMP-001, Sorafinib, Dox in induced pluripotent stem cell cardiomyocytes (iPSc) compared with AMP-001 in MDA-MB-231 triple negative breast cancer cells and F) IC-50 of Dox in iPSc cells.

### 8.0 Discussion

Due to the lack of early detection, human gastric cancer remains as the second most common cause of death from cancer worldwide^32^. Chemotherapies have become inevitable despite high side effects. However, the therapeutic efficacy decreases when cancer cells develop resistance to chemotherapeutics. Combinational chemotherapies have been a mainstay in the treatment of disseminated malignancies for a long time^33-35^ and have been widely used for minimizing acquired resistance, enhancing cell-intrinsic drug synergy with chemotherapy or for maximizing cumulative drug dose^36a^. One of the main reasons for tumor cells to develop resistance is downregulation of cell death pathways by cancer cells to apoptosis. New methods are required to activate these dysregulated cell death pathways and to sensitize the low-responsive resistant tumor cells which are otherwise non-responsive to any therapy.

The present study proposes targeting the chemotherapeutics using a tumor specific biomarker Cathespin B. Cathepsin B is overexpressed in many cancers including gastric cancer making it a unique biomarker of cancer to which drugs can be targeted^21^. Valine-citrulline dipeptide is one such linker which is cleavable by Cathespsin B^20^. The drug design AMP-001 has to take into consideration the release of apoptogen (apoptosis inducing ligand) at tumor sites in order to be selective to cancer cells. The pegylation of the basic apoptogen core is expected to increase water solubility and make it stable in the circulatory blood for a long time to reach the tumor^37^. The parent molecule α-tocopheryl succinate (α-TOS), although showed high promise in both *in vitro* and *in vivo*, low solubility in water and a serious toxicity in immunocompetent animal model limited its utility^23^. This observation could explain the lack of clinical product so far for α-TOS. This prompted us to redesign apoptogen keeping in mind with a targeting moiety, selectivity and high solubility in water.

Most clinically used chemotherapeutics are not selective to cancer. Moreover, these drugs do not address the fundamental aberrations of cancer cells ability to circumvent interventions by dysregulating cell death pathways and desensitize themselves to intervention, irrespective of nature and/or mode of intervention. Hence, our first approach was to sensitize the low or non-responsive tumor cells by activating CD95 pathway, one of the major dysregulated cell death pathways. The second approach was to take advantage of activation technology to ascertain if it entitles synergy with the standard chemotherapeutics. The significant decrease of IC_50_ for the combination of doxorubicin and AMP-001 for gastric cancer cells (Fig 3-4) reveals that lower dose of doxorubicin could be clinically useful if AMP-001 is used as neoadjuvant to chemotherapy. The tumor cells were pretreated for 6-12 hrs. depending upon cancer cell type and allow the activation of CD95 before we administer chemotherapy. Several vitamin E (VE) analogues, such as α-TOS, α-TOH and VES have been documented as combinations with chemotherapeutics, such as doxorubicin, statins^38^, gefitinib^39^, celecoxib^40^, SU11274^41^, GW966^42^, oridonin^43^ and baicalein^44^ for potential synergetic effects. However, our leading compound AMP-001 is different from the others in terms of ability to target cancer cells sparing normal cells. Activation of cell death pathways in normal cells, for example in brain leads to neuro degenerative diseases^13^ and hence, targeting is essential.

Our preclinical studies *in vitro* and *in vivo* revealed that AMP-001 has dual effects; at low dose it acts as a tumor sensitizer inducting a reasonable cell death while, at higher doses it shows anti-tumorigenicity on its own with little or no observable toxicity in non-target organs^12^. However, for our current studies we used AMP-001 as a tumor sensitizer. The main objective is to enhance the efficacy of the currently used chemotherapeutics using AMP-001 as a neoadjuvant to chemotherapy. The evidence showed that AMP-001 synergized with doxorubicin and thereby, circumvents the cells which are resistant to doxorubicin in human gastric cancer cells. Since doxorubicin is a first-line chemotherapeutic agent in clinical cancer treatments but limited in its use due its cardiotoxicity, the combination of doxorubicin and AMP-001 may be more effective in the clinical setting compared to doxorubicin alone in order to reduce the off-target toxicity and to overcome cancer drug resistance.

In order for a potential clinical translation of AAAPT technology, we have studied the tumor regression *in vivo* using triple negative breast cancer xenograft model so that the synergistic potential of AMP-001 could be extendable to *in vivo* situations. Doxorubicin has been combined with many other drugs for a potential synergistic effects^19^. However, AMP-001 has a unique ability to sensitize tumor cells at low dose, while inducing tumor regression at higher doses. In our alternate studies, we have shown that at 200mg/Kg dose in a triple negative breast cancer xenograft model tumor regression was more than 80 percent compared to control^12^(supplementary data Fig. 1) Here, in this study we have shown that the combination of AMP-001 and doxorubicin showed a synergistic tumor regression (p < 0.005) compared to individual drugs (Fig 7). We have fixed AMP-001 at a lower dose and used it as a sensitizing agent rather than as an anti-tumor agent. The combination treatment also showed non-significant changes in weight loss suggesting a favorable toxicity (safety) profile for the combination. Conventional doxorubicin treatment, despite being anti-tumorigenic riddles with cardiotoxicity issues. Hence, we conducted studies on the relative toxicity profile of AMP-001 and doxorubicin in human induced pluripotent stem cell cardiomyocytes (iPSCs) using a proprietary assay developed by Ionic Transport Assays Inc. Adult human induced pluripotent stem cell-derived (hIPSC) cardiomyocyte technology, such as iCell ^R^ Cardiomyocytes, offers the opportunity to accelerate the development of new therapeutic agents by providing a relevant human target for efficacy and safety without going through the costly animal studies. Our results show that the conventional chemotherapy such as doxorubicin and sorafinib showed 9.9 μM and 40 μM respectively while, AMP-001 was greater than 250 μM (Fig 9F and E) indicating a better safety profile for AMP-001. It is to be noted that AMP-001 showed lower IC-50 (32 μM) for TNBC cells indicating that the efficacy is as good as many FDA approved chemotherapeutics. The IC-50 values corroborated well with the photomicrograph of cardiomyocytes treated with doxorubicin showing significant cell death compared to control DMSO and 100 μMAMP-001 (Fig 10 C Vs A and D).

The possible mechanism of AMP-001 synergetic effect was also explored. Combination of AMP-001 and doxorubicin induced increased intracellular ROS level, determined by the shift of DCFCA fluorescence peak to right (Fig 5A-B). The selectivity of AMP-001 to cancer is due to the targeting vector valine-citrulline dipeptide which is cleaved by tumor specific Cathepsin B. However, recent studies on ROS mediated damage to cells also indicates, the kinetics of repair is almost twice faster in normal cells compared to cancer cells^46^. Thus, enhancing ROS selectively in cancers could be a better strategy since normal cells gets repaired faster than cancer cells and targeting helps minimizing affecting normal cells. Previous studies have demonstrated how vitamin E analogues generate ROS species which led to cell apoptosis. One possibility is that vitamin E analogues are known to interact with the UbQ-binding sites resulting in destroying mitochondrial electron doxorubicin chain^45-48^. The shift of red fluorescence to green fluorescence for combined doxorubicin and AMP-001 (Fig 6B) indicates the significant reduction in the mitochondrial potential making mitochondria in cancer cells as a potential target.

In our studies, we found combinational drugs seem to affect mitochondria membrane potential and triggered Bcl_2_/Bax dependent mitochondria apoptotic cascade more effectively than the mono-drug treatment. With the elevated cell stress induced by drugs, Mitogen Activated Protein Kinase (MAPK) signaling pathway was also activated (Fig 7).

MAPKs represent a family of kinases that transduce diverse extracellular stimuli including proapoptotic agents) to the nucleus via kinase cascades to regulate proliferation, DNA synthesis arrest, differentiation and apoptosis phenomena. MAPKs are activated through phosphorylation of specific threonines and tyrosines by dual specificity kinases via a four-step kinase cascade. There are three well-defined MAPK pathways in mammalian cells: the ERK1/ERK2 cascade and the stress-activated JNK MAPK cascades. Our studies document that either AMP-001 or doxorubicin triggered apoptosis involves ERK1, MEK1, and JNK1. Both ERK1/2 and JNK1 were activated after drug treatment and were documented by detection of the active (phosphorylated) forms of these kinases using antibodies specific for the active enzyme. However, the combination of AMP-001 and doxorubicin yielded a stronger band in Western Blotting which is corroborated well with the other *in vitro* data such as reduction in IC-50, enhancement of ROS and disintegration of mitochondrial permeability. In other words, these data prove that AMP-001 is synergistic with doxorubicin to enhance its efficacy.

In essence, multiple pathways are involved for the synergistic effect of AMP-001 with doxorubicin. It is important to note that the dysregulation of cell death pathways is manifested in many cancers including gastric cancer through the downregulation of CD95^3^. The appearance of band in Western Blotting at 43 KDa seems to explain the activation of CD95 pathway which has been shown to sensitize tumor cells^31^. Hence, activation of CD95 pathway selectively is presumed to sensitize tumor cells. Studies on cis-platin in testicular cancer, for example concludes that loss of activation of CD95 pathway resulted in developing resistance to cis-platin^4^. Further studies are necessary to confirm this hypothesis.

We can only speculate the potential mechanism of action based on our limited studies reported in our patent disclosure which includes a) release of AIF1 from mitochondria Fig (2 Supplementary), b) reduction of downstream Bcl2/Bax ratio (Fig 3A-B, Supplementary data), NF-kB inhibition (Fig 4-5, supplementary data), PARP cleavage (Fig 6, supplementary data) and generation of reactive oxygen species (Fig 7, ROS, data). Based on the limited data, it could be hypothesized that both nucleus and mitochondria may be involved. ROS, generated by AMP-001/002 is presumably translocated Bax into mitochondria (data not shown) triggering NF-kB inhibition which, in turn induces cell death presumably through α-TNF related apoptosis inducing ligand (TRAIL) or interferons (IFNs). Similarly, release of apoptosis inducing factor (AIF-1) from mitochondria to cytosol and subsequent reduction of downstream Bcl2/Bax proteins triggers irreversible cell death through caspase independent pathway. On the other hand, cell death CD95 activation is expected to mediate cell death through FADD/procaspase and caspase 3/7 pathway. Cleavage of DNA repair enzyme PARP augments cell death and established to sensitize both CSCs and resistant tumor cells. This contrasts with doxorubicin which is shown to hyperactivate NF-kB and PARP and impairs mitochondrial respiratory chain complex I leading to cell death in cardiomyocytes. Thus, inhibition of NF-kB pathway and PARP cleavage protects cardiomyocytes reducing the potential cardiotoxicity. A potential mechanism of action is described in Fig 10.

**Fig 10:**
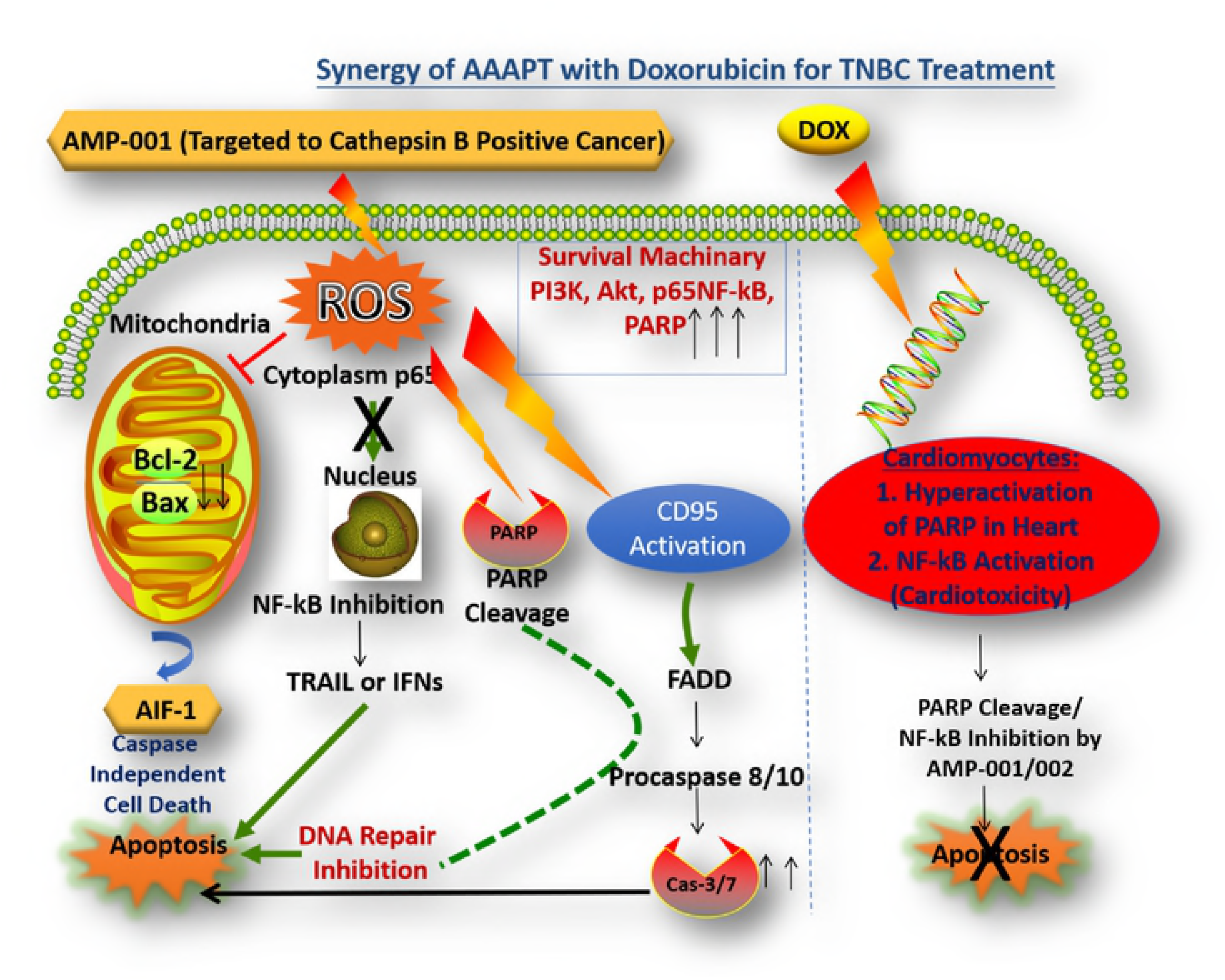
Potential mechanism of AMP-001 mediating synergy with Doxorubicin.

## Conclusions

Since cancer is a group of diseases, several targets and pathways are involved in the progression of the disease. Although, conventional drugs apart from nonspecific chemotherapeutics target specific target (s), each drug works with specific targets/pathways. The combination of AAAPT leading molecule with a conventional chemotherapy brings multiple pathways involved in cancer cells desensitization process.

In summary, the effects of novel vitamin E analogue AMP-001 is as a preclinical candidate is described. AMP-001 enhanced doxorubicin induced apoptosis in human gastric cancer cells via ROS-dependent mitochondrial apoptotic pathway and MAPK pathway. The efficacy of doxorubicin antitumor activity can be amplified by combining it with AMP-001, which permits lower dose use in patients without affecting the positive clinical outcome. These effects are confirmed both *in vitro* and *in vivo*. Our current studies indicated that AMP-001 might be a potential candidate in synergetic with the existing FDA approved chemotherapeutic drugs for treatment of gastric cancer. Ultimately, the combination treatments success depends on how low dose can be made effective in a clinical situation, reduced overall toxicity and improved patient compliance. Further studies are planned to optimize the dose combination as to which one has higher efficacy and low toxicity compared to the existing treatments.

## Acknowledgements

Part of this research was supported by the grant SBIR NIH R43CA183353.

